# C-section and systemic inflammation synergize to disrupt the neonatal gut microbiota and brain development in a model of prematurity

**DOI:** 10.1101/2023.10.20.563256

**Authors:** Cécile Morin, Flora Faure, David Guenoun, Irvin Sautet, Sihao Diao, Valérie Faivre, Jennifer Hua, Leslie Schwendimann, Amazigh Mokhtari, Rebeca Martin, Sead Chadi, Charlie Demené, Andrée Delahaye-Duriez, Rochellys Diaz-Heijtz, Bobbi Fleiss, Boris Matrot, Sandrine Auger, Mickael Tanter, Juliette Van Steenwinckel, Pierre Gressens, Cindy Bokobza

**Author notes:** Corresponding author: Dr Cindy Bokobza, Inserm Unité 1141, Hôpital Robert Debré 48 Boulevard Sérurier, 75019 Paris, France <u>.:</u> <u>Mail:</u>. Equal contribution (Juliette Van Steenwinckel, Pierre Gressens, Cindy Bokobza). **Author Contributions:** CM, FF, VF, JH, LS, RM, CD, RDH, BF, BM, SA, MT, JVS, PG, and CB: study concept and design. CM, FF, DG, IS, SD, VF, JH, LS, AM, RM, CD, ADD, RDH, BF, BM, SA, MT, JSV, PG, and CB: data acquisition and analysis. CM, FF, VF, JH, AM, CD, ADD, RDH, BF, BM, SA, MT, JVS, PG, and CB: drafting of the text and figures. **Competing Interest Statement:** The authors declare no competing interest.

## Abstract

Infants born very preterm (below 28 weeks of gestation) are at high risk of developing neurodevelopmental disorders, such as intellectual deficiency, autism spectrum disorders, and attention deficit. Preterm birth often occurs in the context of perinatal systemic inflammation due to chorioamnionitis and postnatal sepsis (Dammann, O. and Leviton, A., *Intermittent or sustained systemic inflammation and the preterm brain*. Pediatr Res, 2014. **75**(3): p. 376-80). In addition, C-section is often performed for very preterm neonates to avoid hypoxia during a vaginal delivery (Luca, A.,et al., *Birth trauma in preterm spontaneous vaginal and cesarean section deliveries: A 10-years retrospective study.* PloS one,2022, 17(10), e0275726.) We have developed and characterized a mouse model based on intraperitoneal injections of IL-1β between postnatal days one and five to reproduce perinatal systemic inflammation (Favrais, G.,et al., *Systemic inflammation disrupts the developmental program of white matter.* Ann Neurol,2011. **70**(4): p. 550-65). This model replicates several neuropathological, brain imaging, and behavioral deficits observed in preterm infants. We hypothesized that C-sections could synergize with systemic inflammation to induce more severe brain abnormalities. We observed that C-sections significantly exacerbated the deleterious effects of IL-1β on reduced gut microbial diversity, increased levels of circulating peptidoglycans, abnormal microglia/macrophage reactivity, impaired myelination, and reduced functional connectivity in the brain relative to vaginal delivery plus intraperitoneal saline. These data demonstrate the deleterious synergistic effects of C-section and neonatal systemic inflammation on brain maldevelopment and malfunction, two conditions frequently observed in very preterm infants, who are at high risk of developing neurodevelopmental disorders.

**Significance Statement:** In a well-established mouse model of the encephalopathy of prematurity, we observed that C-section exacerbates the deleterious effects of neonatal systemic inflammation (intraperitoneal injections of IL-1β between postnatal days one and five) on reduced gut microbial diversity, increased levels of circulating peptidoglycans, abnormal microglia/macrophage reactivity, impaired myelination, and reduced brain functional connectivity. These data demonstrate the deleterious synergistic effects of C-section and neonatal systemic inflammation, two conditions frequently observed in very preterm infants, who are at high risk of developing neurodevelopmental disorders.

## Introduction

Preterm delivery (WHO definition: below 38 weeks of gestation) occurs in more than 10% of births worldwide and is associated with a very significant increased risk of motor deficits (cerebral palsy), intellectual deficiency, and psychiatric disorders (including autism spectrum disorder and attention deficit) relative to term delivery [1–3]

Imaging and post-mortem studies of preterm infants combined with experimental models have identified the brain abnormalities associated with a poor long-term neurological prognosis. These abnormalities, grouped under the term encephalopathy of prematurity (EoP), include diffuse white matter injury related to delayed myelination induced by the blockade of maturation of oligodendrocyte progenitor cells (OPCs), microglia and astrocyte activation, which play a pivotal role in the pathogenic cascade, a reduction in the density of subsets of interneurons, the abnormal microstructure of cortical grey matter, and reduced brain connectivity [4–9].

Epidemiological and clinical studies have highlighted an important role for sustained systemic inflammation in the etiology of EoP [10]. Several conditions can contribute to perinatal inflammation in preterm infants, including prenatal chorioamnionitis, postnatal sepsis, mechanical ventilation, and delayed necrotizing enterocolitis [11, 12].

Gut microbiota dysbiosis has been shown to be associated with several neurodevelopmental disorders (NDDs), including autism spectrum disorder, although its pathogenic role is still a subject of debate [13–16]. In the context of preterm delivery, several studies have reported a reduced diversity of gut microbiota that can persist for several weeks or months after birth [17–21]. Several factors have been shown to potentially induce or be associated with dysbiosis in preterm infants, including cesarean delivery (C-section), preventing colonization by the vaginal microbiota, exposure to multiple rounds of broad-spectrum antibiotics, sustained systemic inflammation, and delayed introduction of oral feeding [22–25].

C-sections are often performed for very preterm neonates to avoid the hypoxia that occurs during vaginal delivery. However, several epidemiological studies have shown an association between C-sections and NDDs [24, 26, 27]. Based on the evidence cited above, this raises the question of potential deleterious interactions between C-section and sustained systemic inflammation in the genesis of the EoP and whether these two factors act independently or synergize to worsen the dysbiosis and associated brain maldevelopment that lead to NDDs.

We have developped a rodent model of EoP based on the neonatal intraperitoneal injection of the inflammatory cytokine IL-1β from postnatal day (P) 1 to P5, which recapitulates several key hallmarks of human EoP [16, 28–36]. We have used this model to show that pro-inflammatory activation of microglia/macrophages induced by IL-1β is a major event that leads to the blockade of OPC maturation and the subsequent delay in myelination [29, 32].

Here, we used this EoP model to combine C-section and neonatal systemic inflammation (IL-1β) and determine the impact of each factor or their combination on the gut microbiota, microglia activation, myelination, brain connectivity, and behaviour (Fig. 1A) relative to the control condition of vaginal delivery (V-delivery) and neonatal systemic saline (PBS).

**Figure 1.**
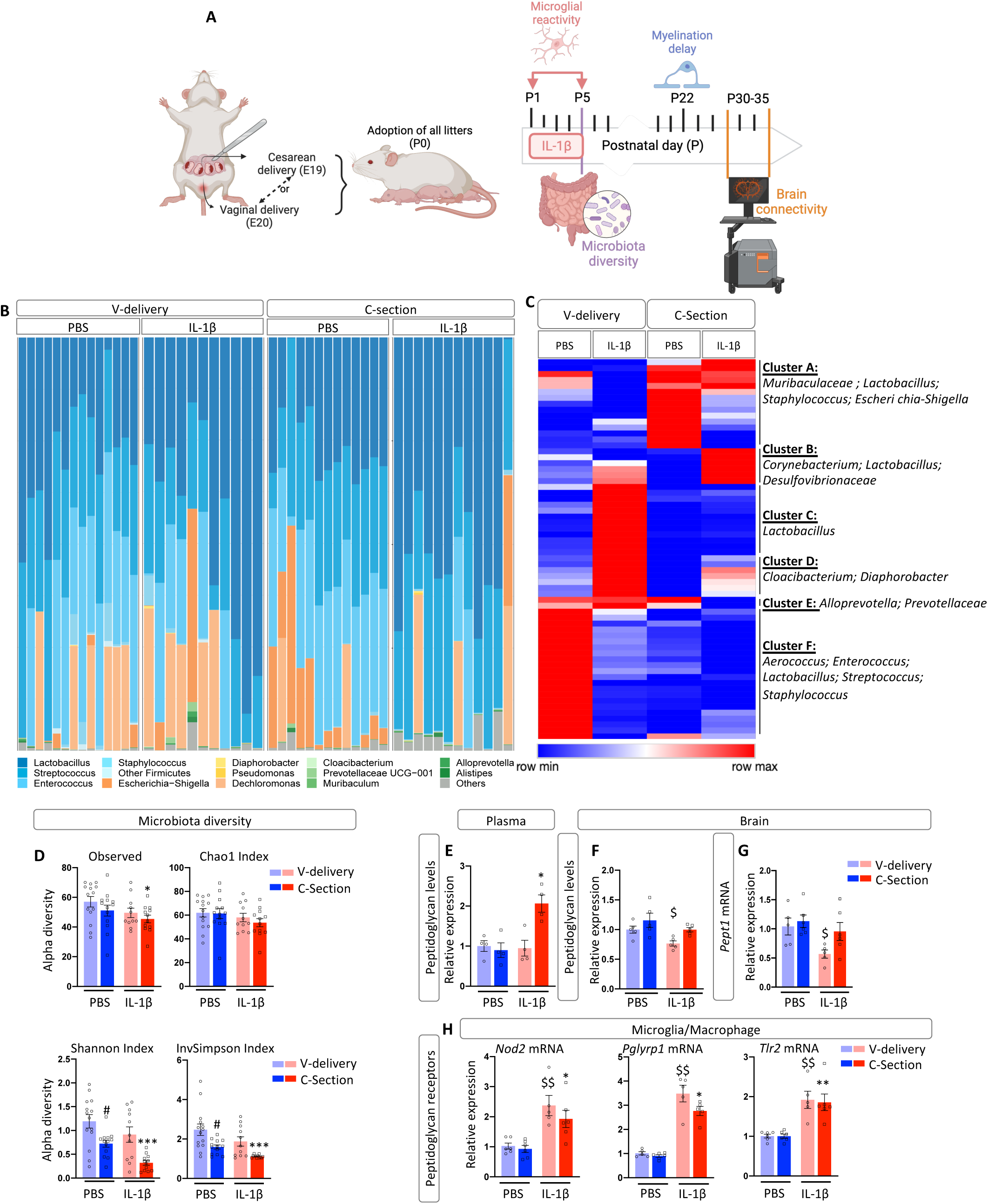
Impact of IL-1β and C-section on microbiota composition and peptidoglycan release. (A) Schematic representation of the experimental protocol. (B) Table of abundance of operational taxonomic units (OTUs). Analysis of the individual distribution of the genera that make up the intestinal microbiota at P5 in V-delivery ± IL-1β or C-section ± IL-1β mice. Each phylum is represented by a different color. (C) Heatmap representation of the differential analysis of the OTUs (BH p<0.05) by hierarchical clustering using Pearson’s correlation. (D) Diversity of the intestinal microbiota at P5 depending on the delivery route and the presence or absence of systemic inflammation. Alpha diversity was determined using the following four indexes: Observed, Chao1, Shannon, and InvSimpson. n=12-14 per group from at least two different cages per group. (E-F) Relative expression of peptidoglycan levels in the plasma (E) and brain (F) at P5 of V-delivery ± IL-1β or C-section ± IL-1β mice. (G) Relative expression of *Pept1* mRNA at P5 in the anterior brain of V-delivery ± IL-1β or C-section ± IL-1β mice. (H) Relative expression of microglial peptidoglycan receptor mRNA at P5 in CD11B+ cells of V-delivery ± IL-1β or C-section ± IL-1β mice. Data are presented as a scatter plot with a bar (mean + SEM). The Kruskal-Wallis Test was used followed by an uncorrected Dunn’s test. All groups were compared to the V-delivery+PBS group.

## Results

### Impact of C-section and systemic inflammation on the gut microbiota and peptidoglycan levels

We studied the composition of the gut microbiota at P5 from colon samples to determine whether C-section and systemic inflammation influence this microbiota. Amplicon sequencing of bacterial 16S-rRNA generated operational taxonomic units (OTUs). Firmicutes, including the genera Lactobacillus, Streptococcus, Enterococcus, and Staphylococcus, was the predominant phylum in the four mouse groups, followed by Proteobacteria and Bacteroidota (Fig. 1B, Supplementary Fig. 1). We used hierarchical clustering analysis to group the OTUs according to their relative abundance within the four groups (Table 1). The effect of C-section and IL-1β-exposure on the abundance of various OTUs was evident, and they were divided into six clusters (A to F) (Fig. 1C). The abundance of the OTUs corresponding to the Lactobacillus genera was highly heterogeneous between groups. Certain Lactobacillus OTUs were found to be over-represented in clusters A and B and under-represented in clusters C and F (Fig. 1C). In addition, cluster F showed the proliferation of Streptococcus and Enterococcus OTUs in the V-delivery+PBS group, whereas these species were less abundant in the other groups, even disappearing in the C-section+IL-1β group (Fig. 1C, Supp. Fig. 1A). Furthermore, OTUs associated with Proteobacteria, specifically those relevant to Escherichia-Shigella in cluster A, were more abundant only in the C-section+PBS group (Fig. 1C, Supplementary Fig. 1B). Finally, IL-1β-exposure specifically promoted over-representation of OTUs, such as Corynebacterium and Desulfovibrionaceae (Cluster B), which could be modulated by C-section, as seen with Cloacibacterium and Diaphorobacter in cluster D (Fig. 1C).

**Table 1.**

| Cluster A |  |
| --- | --- |
| Cluster 27 | Bacteria;Bacteroidota;Bacteroidia;Bacteroidales;Muribaculaceae;unknown genus;unknown species |
| Cluster 110 | Bacteria;Bacteroidota;Bacteroidia;Bacteroidales;Muribaculaceae;unknown genus;unknown species |
| Cluster 117 | Bacteria;Bacteroidota;Bacteroidia;Bacteroidales;Muribaculaceae;unknown genus;unknown species |
| Cluster 1 | Bacteria;Firmicutes;Bacilli;Lactobacillales;Lactobacillaceae;Lactobacillus;Multi-affiliation |
| Cluster 141 | Bacteria;Firmicutes;Bacilli;Lactobacillales;Lactobacillaceae;Lactobacillus;Multi-affiliation |
| Cluster 126 | Bacteria;Firmicutes;Bacilli;Lactobacillales;Lactobacillaceae;Lactobacillus;Multi-affiliation |
| Cluster 135 | Bacteria;Firmicutes;Bacilli;Lactobacillales;Lactobacillaceae;Lactobacillus;Multi-affiliation |
| Cluster 36 | Bacteria;Firmicutes;Bacilli;Lactobacillales;Lactobacillaceae;Lactobacillus;Multi-affiliation |
| Cluster 72 | Bacteria;Firmicutes;Bacilli;Lactobacillales;Lactobacillaceae;Lactobacillus;Multi-affiliation |
| Cluster 94 | Bacteria;Firmicutes;Bacilli;Lactobacillales;Lactobacillaceae;Lactobacillus;Multi-affiliation |
| Cluster 198 | Bacteria;Firmicutes;Bacilli;Lactobacillales;Lactobacillaceae;Lactobacillus;unknown species |
| Cluster 21 | Bacteria;Firmicutes;Bacilli;Staphylococcales;Staphylococcaceae;Staphylococcus;Multi-affiliation |
| Cluster 165 | Bacteria;Firmicutes;Bacilli;Staphylococcales;Staphylococcaceae;Staphylococcus;Multi-affiliation |
| Cluster 115 | Bacteria;Proteobacteria;Gammaproteobacteria;Enterobacterales;Enterobacteriaceae;Escherichia-Shigella;Multi-affiliation |
| Cluster 2 | Bacteria;Proteobacteria;Gammaproteobacteria;Enterobacterales;Enterobacteriaceae;Escherichia-Shigella;Multi-affiliation |

| Cluster B |  |
| --- | --- |
| Cluster 140 | Bacteria;Actinobacteriota;Actinobacteria;Corynebacteriales;Corynebacteriaceae;Corynebacterium;unknown species |
| Cluster 128 | Bacteria;Desulfobacterota;Desulfovibrionia;Desulfovibrionales;Desulfovibrionaceae;unknown genus;unknown species |
| Cluster 24 | Bacteria;Firmicutes;Bacilli;Lactobacillales;Lactobacillaceae;Lactobacillus;unknown species |
| Cluster 119 | None |
| Cluster 143 | None |
| Cluster 99 | None |

| Cluster C |  |
| --- | --- |
| Cluster 122 | Bacteria;Firmicutes;Bacilli;Lactobacillales;Lactobacillaceae;Lactobacillus;Multi-affiliation |
| Cluster 118 | Bacteria;Firmicutes;Bacilli;Lactobacillales;Lactobacillaceae;Lactobacillus;Multi-affiliation |
| Cluster 11 | Bacteria;Firmicutes;Bacilli;Lactobacillales;Lactobacillaceae;Lactobacillus;Multi-affiliation |
| Cluster 136 | Bacteria;Firmicutes;Bacilli;Lactobacillales;Lactobacillaceae;Lactobacillus;Multi-affiliation |
| Cluster 43 | Bacteria;Firmicutes;Bacilli;Lactobacillales;Lactobacillaceae;Lactobacillus;Multi-affiliation |
| Cluster 15 | Bacteria;Firmicutes;Bacilli;Lactobacillales;Lactobacillaceae;Lactobacillus;Multi-affiliation |
| Cluster 4 | Bacteria;Firmicutes;Bacilli;Lactobacillales;Lactobacillaceae;Lactobacillus;Multi-affiliation |
| Cluster 106 | Bacteria;Firmicutes;Bacilli;Lactobacillales;Lactobacillaceae;Lactobacillus;unknown species |
| Cluster 138 | Bacteria;Firmicutes;Bacilli;Lactobacillales;Lactobacillaceae;Lactobacillus;unknown species |
| Cluster 12 | Bacteria;Firmicutes;Bacilli;Lactobacillales;Lactobacillaceae;Lactobacillus;unknown species |
| Cluster 35 | Bacteria;Firmicutes;Bacilli;Lactobacillales;Lactobacillaceae;Lactobacillus;unknown species |
| Cluster 18 | Bacteria;Firmicutes;Bacilli;Lactobacillales;Lactobacillaceae;Lactobacillus;unknown species |
| Cluster 34 | None |
| Cluster 47 | None |

| Cluster D |  |
| --- | --- |
| Cluster 60 | Bacteria;Bacteroidota;Bacteroidia;Flavobacteriales;Weeksellaceae;Cloacibacterium;Multi-affiliation |
| Cluster 30 | Bacteria;Proteobacteria;Gammaproteobacteria;Burkholderiales;Comamonadaceae;Diaphorobacter;Multi-affiliation |
| Cluster 127 | None |
| Cluster 81 | None |
| Cluster 61 | None |

| Cluster E |  |
| --- | --- |
| Cluster 116 | Bacteria;Bacteroidota;Bacteroidia;Bacteroidales;Prevotellaceae;Alloprevotella;unknown species |
| Cluster 85 | Bacteria;Bacteroidota;Bacteroidia;Bacteroidales;Prevotellaceae;Prevotellaceae UCG-001;unknown species |

| Cluster F |  |
| --- | --- |
| Cluster 32 | Bacteria;Firmicutes;Bacilli;Lactobacillales;Aerococcaceae;Aerococcus;Multi-affiliation |
| Cluster 17 | Bacteria;Firmicutes;Bacilli;Lactobacillales;Enterococcaceae;Enterococcus;Enterococcus faecalis |
| Cluster 5 | Bacteria;Firmicutes;Bacilli;Lactobacillales;Enterococcaceae;Enterococcus;Multi-affiliation |
| Cluster 55 | Bacteria;Firmicutes;Bacilli;Lactobacillales;Enterococcaceae;Enterococcus;Multi-affiliation |
| Cluster 8 | Bacteria;Firmicutes;Bacilli;Lactobacillales;Enterococcaceae;Enterococcus;Multi-affiliation |
| Cluster 10 | Bacteria;Firmicutes;Bacilli;Lactobacillales;Lactobacillaceae;Lactobacillus;Multi-affiliation |
| Cluster 20 | Bacteria;Firmicutes;Bacilli;Lactobacillales;Lactobacillaceae;Lactobacillus;Multi-affiliation |
| Cluster 26 | Bacteria;Firmicutes;Bacilli;Lactobacillales;Lactobacillaceae;Lactobacillus;Multi-affiliation |
| Cluster 29 | Bacteria;Firmicutes;Bacilli;Lactobacillales;Lactobacillaceae;Lactobacillus;Multi-affiliation |
| Cluster 33 | Bacteria;Firmicutes;Bacilli;Lactobacillales;Lactobacillaceae;Lactobacillus;Multi-affiliation |
| Cluster 37 | Bacteria;Firmicutes;Bacilli;Lactobacillales;Lactobacillaceae;Lactobacillus;Multi-affiliation |
| Cluster 57 | Bacteria;Firmicutes;Bacilli;Lactobacillales;Lactobacillaceae;Lactobacillus;Multi-affiliation |
| Cluster 87 | Bacteria;Firmicutes;Bacilli;Lactobacillales;Lactobacillaceae;Lactobacillus;Multi-affiliation |
| Cluster 28 | Bacteria;Firmicutes;Bacilli;Lactobacillales;Lactobacillaceae;Lactobacillus;unknown species |
| Cluster 44 | Bacteria;Firmicutes;Bacilli;Lactobacillales;Lactobacillaceae;Lactobacillus;unknown species |
| Cluster 13 | Bacteria;Firmicutes;Bacilli;Lactobacillales;Multi-affiliation;Multi-affiliation;Multi-affiliation |
| Cluster 3 | Bacteria;Firmicutes;Bacilli;Lactobacillales;Streptococcaceae;Streptococcus;unknown species |
| Cluster 39 | Bacteria;Firmicutes;Bacilli;Lactobacillales;Streptococcaceae;Streptococcus;unknown species |
| Cluster 45 | Bacteria;Firmicutes;Bacilli;Lactobacillales;Streptococcaceae;Streptococcus;unknown species |
| Cluster 90 | Bacteria;Firmicutes;Bacilli;Lactobacillales;Streptococcaceae;Streptococcus;unknown species |
| Cluster 102 | Bacteria;Firmicutes;Bacilli;Staphylococcales;Staphylococcaceae;Staphylococcus;Multi-affiliation |
| Cluster 6 | Bacteria;Firmicutes;Bacilli;Staphylococcales;Staphylococcaceae;Staphylococcus;Multi-affiliation |

We further evaluated the α-diversity of the gut microbiota using various indexes. The observed species and Chao1 index were used to determine the number of species present, while the Shannon and InvSimpson indexes assessed the variety and evenness of the species [37]. The C-section+IL-1β group showed a significant decrease in observed diversity (p=0.017). In addition, we observed a significant decrease in the Shannon and InvSimpson indexes in the C-section+PBS group (p=0.04). This effect was even more pronounced in the C-section+IL-1β group (p<0.0001) relative to the V-delivery+PBS group (Fig.1D).

Peptidoglycan (PGN) motifs derived from commensal gut microbiota can be translocated into the developing brain and sensed by specific pattern recognition receptors (PRRs) of the innate immune system [38, 39]. We evaluated PGN levels in the plasma and brain at P5 [40, 41]. Plasma PGN levels in the C-section+IL-1β group were significantly higher than those in the V-delivery+PBS group (p=0.039) (Fig. 1E). By contrast, brain PGN levels in the V-delivery+IL-1β group were significantly lower than those in the V-delivery+PBS group (p=0.046) (Fig.1F). This reduction in brain PGN levels could be linked to the significant under-expression (p=0.036) of brain *Pept1* mRNA, one of the PGN transporters within the brain, in the V-delivery+IL-1β group (Fig. 1G).

PGNs interact with two families of specific pattern recognition receptors (PRRs): PGN-recognition proteins (PGLYRP1-4) and NOD-like receptors (NOD1-2) [42, 43]. In a previous study, we performed transcriptomic analysis (mRNA microarrays) of CD11B^+^ brain cells from V-delivery+PBS and V-delivery+IL-1β groups using the same model of EoP. These data showed higher *Nod1-2 and Tlr2* and only *Pglyrp1* mRNA expression in CD11B^+^ microglia/macrophages in the V-delivery+IL-1β group (Supplementary Fig.2). Here, we extended this analysis to the four studied groups by qRT-PCR analysis of CD11B^+^ cells (Fig. 1H). We confirmed the significant overexpression of *Nod2* (p=0.004), *Pglyrp1* (p=0.004), and *Tlr2* (p=0.003) mRNA in the V-delivery+IL-1β group and observed comparable overexpression of these three mRNAs the C-section+IL-1β group (p=0.024, 0.047, and 0.006, respectively).

### Impact of C-section and systemic inflammation on microglial activation

We previously showed that CD11B^+^ microglia/macrophages in the V-delivery+IL-1β group overexpress pro-inflammatory markers as soon as two hours after the first IL-1β injection relative to the V-delivery+PBS group [31, 32]. Here, we extended this analysis to the four studied groups by qRT-PCR quantification of several markers of microglia/macrophage reactivity on CD11B^+^ cells (Fig. 2A). We confirmed the significant overexpression at P1 of *prostaglandin-endoperoxide synthase 2 (Ptgs2*, a pro-inflammatory factor)*, tumor necrosis factor* (*Tnf,* a pro-inflammatory factor)*, interleukin 1 receptor antagonist (Il1ra*, an immunomodulatory factor), and *suppressor of cytokine signaling 3 (Socs3*, an immunomodulatory factor) mRNA (p=0.0001, 0.008, and 0.0001, respectively) combined with a delayed increase at P5 of *Arg1* (*arginase 1*, an anti-inflammatory factor) mRNA expression (p=0.03) in the V-delivery+IL-1β group relative to the V-delivery+PBS group. We did not observe any significant difference in mRNA expression between the C-section+PBS and V-delivery+PBS groups. By contrast, at P1, there was very robust induction of *cluster of differentiation 32, (Cd32, a pro-inflammatory factor*)*, Ptgs2, Tnf, Il1ra*, *Socs3*, and *interleukin 4 receptor antagonist*, (*Il-4ra,* an immunomodulatory factor) mRNA expression (p<0.0001) combined with a delayed increase at P5 of *Arg1* mRNA expression (p<0.0001) in the C-section+IL-1β group relative to the V-delivery+PBS group; the overexpression of mRNA was broader and stronger in the C-section+IL-1β group than the V-delivery+IL-1β group.

**Figure 2.**
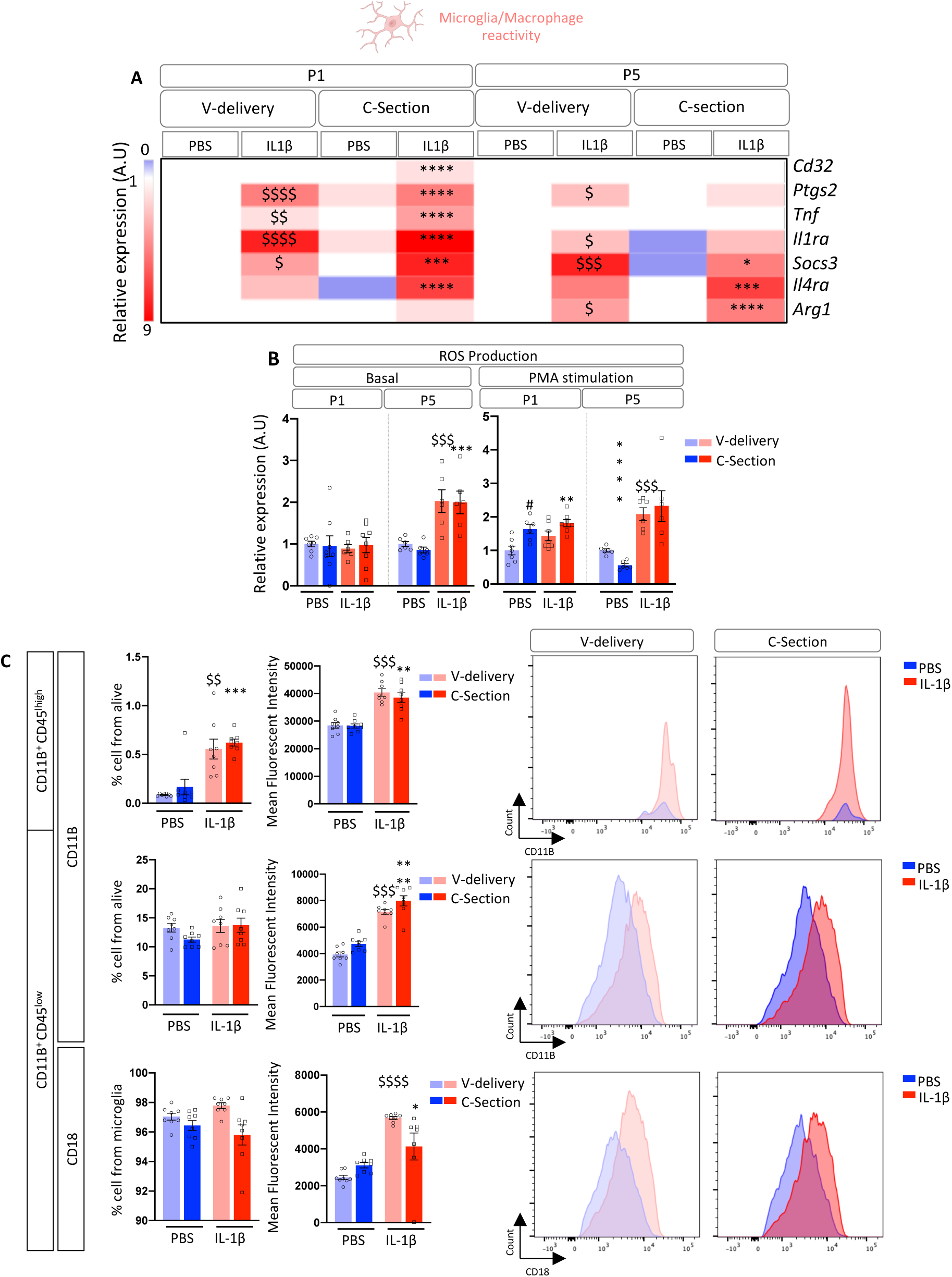
Impact of IL-1β and C-section on microglial reactivity. (A) Heatmap representation of the relative expression of microglial reactivity markers (pro-inflammatory, immune-regulatory, and anti-inflammatory) evaluated by qPCR from CD11B^+^ cells from P1 and P5 brains from V-delivery ± IL-1β or C-section ± IL-1β mice. n=6-9 per group and comparisons between groups and ages were performed using two-way ANOVA followed by a post hoc uncorrected Fisher’s LSD test. All groups were compared to the V-delivery+PBS group. (B) ROS production from CD11B^+^ cells from P1 and P5 brains from V-delivery ± IL-1β or C-section ± IL-1β mice. n= 6-7 per group and comparisons between groups and ages were performed using 2-way ANOVA followed by a post hoc uncorrected Fisher’s LSD test. All groups were compared to the V-delivery+PBS group. (C) Flow cytometry analysis of microglial reactivity in P5 brains from V-delivery ± IL-1β or C-section ± IL-1β mice. Macrophages were defined as CD11B+/CD45^high^. Microglia were defined as CD11B+/CD45^low^. The proportion of microglia and the expression of CD11B and CD18 were evaluated. Data are represented as a scatter plot with a bar (mean + SEM). The Kruskal-Wallis Test was used followed by an uncorrected Dunn’s test, n=8/group. All groups were compared to the V-delivery+PBS group.

We further characterized the microglial/macrophage profile by analyzing the release of reactive oxygen species (ROS) by CD11B^+^ cells sorted at P1 and P5. ROS were measured under basal conditions and after PMA stimulation of sorted CD11B^+^ cells (Fig. 2B) [28, 29]. Under basal conditions, we did not observe any ROS release from P1 CD11B^+^ cells in any experimental group, whereas P5 CD11B^+^ cells sorted from the V-delivery+IL-1β and C-section+IL-1β groups produced a significantly higher amount of ROS (p=0.006 and 0.0008, respectively) than those from the V-delivery+PBS group. Under PMA stimulation, P1 CD11B^+^ cells sorted from the C-section+PBS and C-section+IL-1β groups produced a significantly higher amount of ROS (p=0.025 and 0.0031, respectively) than those of the V-delivery+PBS group. The data for sorted CD11B^+^ cells at P5 under PMA stimulation and basal conditions were comparable.

We further characterized the microglia and infiltrating macrophages in the present model using a previously published gating strategy [44] by quantifying the microglia, defined as CD11B^+^CD45^low^ cells, and infiltrating macrophages, defined as CD11B^+^CD45^high^ cells, at P5 (Fig. 2C). In addition, we also measured the CD11B and CD18 mean fluorescence intensities (MFIs), used as markers of microglia/macrophage activation, at P5 (Fig. 2C). There was a significantly higher proportion of infiltrating CD11B^+^CD45^high^ cells in the V-delivery+IL-1β group than the V-delivery+PBS group (0.56 ± 0.1% and 0.08 ± 0.004%, respectively, p=0.0022). There was a similar increase in the proportion of infiltrating macrophages in the C-section+IL-1β group (0.62 ± 0.1% and 0.08 ± 0.004%, p=0.0002) relative to the V-delivery+PBS group. The infiltrating macrophages showed a significantly higher CD11B MFI in both the V-delivery+IL-1β and C-section+IL-1β groups than the V-delivery+PBS group (p<0.001). We did not observe any modulation in the percentage of CD11B^+^CD45^low^ cells in either the V-delivery+IL-1β or C-section+IL-1β groups relative to the V-delivery+PBS group. However, the microglia showed a significantly higher CD11B MFI in the V-delivery+IL-1β and C-section+IL-1β groups than the V-delivery+PBS group (p=0.0005 and p<0.0001, respectively). This effect was slightly enhanced in the C-section+IL-1β group relative to the V-delivery+IL-1β group. In addition, the microglia showed a significantly higher CD18 MFI in the V-delivery+IL-1β and C-section+IL-1β groups than the V-delivery+PBS group (p<0.0001 and p=0.014). This effect was slightly less in the C-section+IL-1β group than the V-delivery+IL-1β group.

### Impact of C-section and systemic inflammation on the oligodendrocyte lineage and myelination

We next quantified the PDGFRα^+^, PDGFRα^+^/O4^+^, and O4^+^ cell populations by FACS [44] at P5 (Fig. 3A-B). There were significantly fewer PDGFRα^+^ and PDGFRα^+^/O4^+^ cells (p=0.004 and 0.001, respectively) in the V-delivery+PBS group than the V-delivery+PBS group. There was also a significant and comparable decrease in the number of O4^+^ cells (p<0.001) in the V-delivery+IL-1β and C-section+IL-1β groups. However, the PDGFRα^+^/O4^+^ population was significantly smaller in the C-section+IL-1β (p=0.001), whereas it was not in the V-delivery+IL-1β group. Using immunohistochemistry, we quantified the number of oligodendrocyte transcription factor (OLIG)-2^+^ cells at P5 to assess the entire oligodendrocyte lineage (Fig. 3C). The number of OLIG-2^+^ cells in the C-section+IL-1β group was significantly smaller (p=0.029).

**Figure 3.**
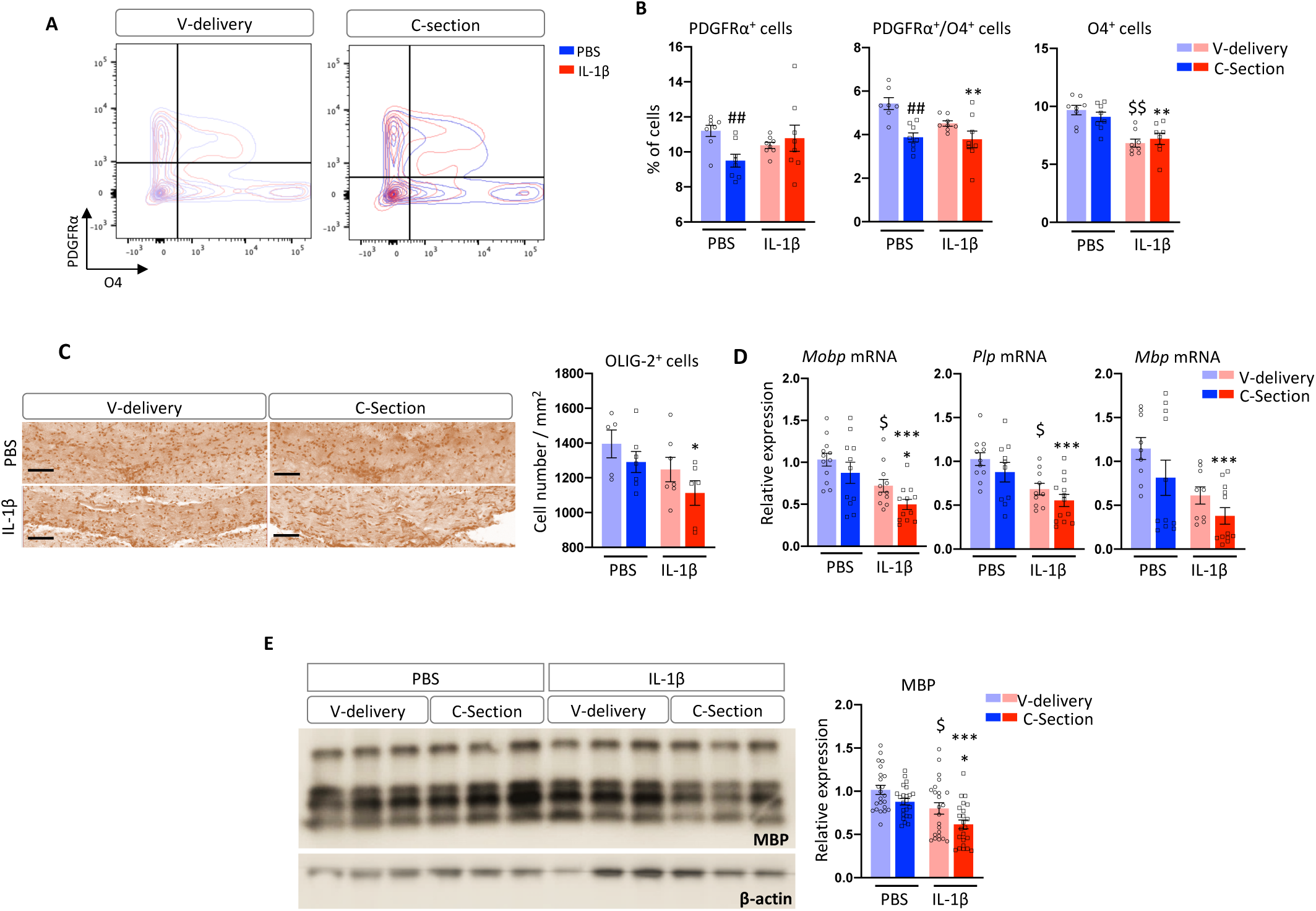
Impact of IL-1β and C-section on white matter injury. (A-B) Flow cytometry analysis of differentiated oligodendrocyte populations (PDGFRα^+^/O4^−^, PDGFRα^+^/O4^+^, and PDGFRα^−^/O4^+^) in P5 brains from V-delivery ± IL-1β or C-section ± IL-1β mice. (C) Total oligodendrocyte lineage cell density by immuno-histochemistry using OLIG-2 labeling in the corpus callosum of P5 V-delivery ± IL-1β or C-section ± IL-1β mice. Representative images were acquired using a Nanozoomer slide scanner. (D) Relative expression of myelin protein mRNA in P22 anterior brains from V-delivery ± IL-1β or C-section ± IL-1β mice evaluated by qPCR. (E) MBP protein quantification by western blotting for V-delivery ± IL-1β or C-section ± IL-1β brains at P22. Data are represented as a scatter plot with a bar (mean + SEM). The Kruskal-Wallis test was used followed by an uncorrected Dunn’s test, n=8/group. All groups were compared to the V-delivery+PBS group.

We then characterized myelination by quantifying the mRNA expression of three myelin proteins in the P22 forebrain by qRT-PCR (Fig. 3D). The expression of *myelin associated oligodendrocyte basic protein (Mobp), proteolipid protein 1 (Plp), and myelin basic protein (Mbp)* mRNA was significantly lower in the V-delivery+IL1β group (p=0.042, 0.014, and 0.06, respectively) and even lower in the C-section+IL1β group (p<0.001).

MBP is among the most abundant myelin proteins in the brain [45]. We previously demonstrated that western blotting evaluation of MBP in the forebrain is a simple and robust method to evaluate the hypomyelination observed in the V-delivery+IL-1β group relative to the V-delivery+PBS group [32, 46]. Here (Fig. 3E), we confirmed that MBP expression is significantly lower in the V-delivery+IL-1β group than the V-delivery+PBS group (p=0.019) and showed such under-expression to be exacerbated in the C-section+IL-1β group (p<0.0001).

### Impact of C-section and systemic inflammation on body weight, behavior, and brain connectivity

Relative to the V-delivery+PBS group, P2 pups in the C-section+PBS group had a significantly higher weight (p=0.0009) and pups in the V-delivery+IL-1β group a significantly lower weight (p=0.002), whereas the weight of the pups in the C-section+IL-1β group was not significantly different (Fig. 4A).

**Figure 4.**
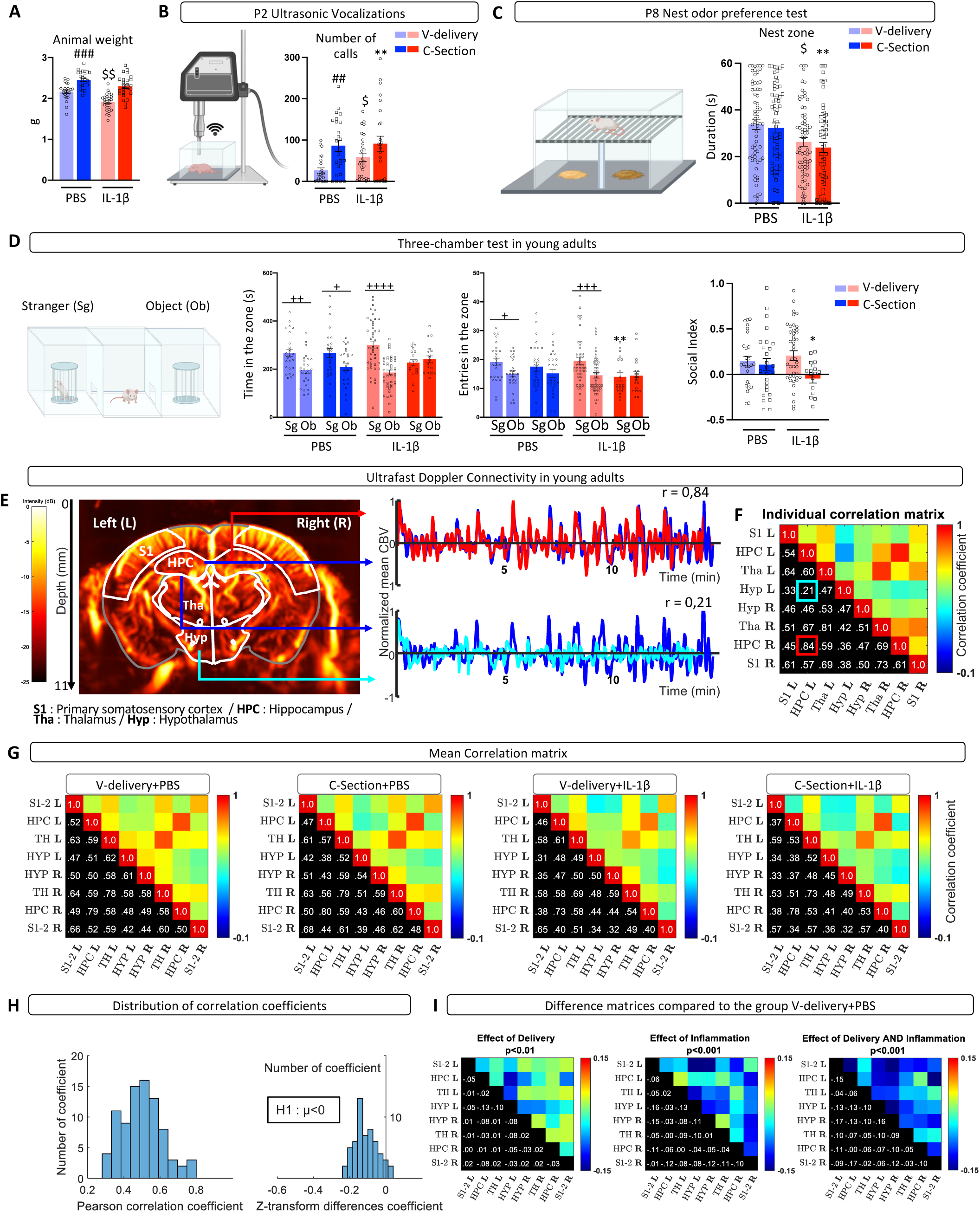
Impact of IL-1β and C-section on weight, behavior, and brain connectivity. (A) Animal weight at P2. (B) Ultrasonic vocalization test evaluating total calls emitted by V-delivery ± IL-1β or C-section ± IL-1β animals at P2 in 3 min. (C) Nest odor preference test at P8 to evaluate maternal recognition. Results are presented as the mean time spent in the nest zone (3 trials of 1 min/animal). (D) Three-chamber test of 5 min at P30-35 comparing V-delivery ± IL-1β or C-section ± IL-1β animals. Data are represented as a scatter plot with a bar (mean + SEM). The Kruskal-Wallis test was used followed by an uncorrected Dunn’s test. All groups were compared to the V-delivery+PBS group. (E) Power Doppler image of a mouse brain in the coronal plane with the ROIs overlayed. fUS were acquired using an Iconeus One ultrasound system, which allowed easy registration on the Allen atlas. Mean CBV signals for the ROIs. From each selected ROI, the CBV signals were spatially averaged and temporally processed to obtain a mean CBV signal to compute the pairwise correlation. The correlation between the right and left hippocampus (top) was high (r=0.84), in contrast to the correlation between the left hippocampus and hypothalamus (down), which was low (r=0.24). (F) Connectivity matrix. The values in the red and blue boxes correspond to the previous mentioned correlations (resp, L/R HPC, and L HPC/Hyp). (G) Mean correlation matrix of each group of interest. (H) Distribution of the correlation coefficients of the mean matrices of pathological groups. Distribution of the difference between the Z-transformed correlation coefficients of the pathological mean matrices and the V-delivery+PBS matrix. From this distribution, it is assumed that the means of the differences between Z-transformed coefficients are smaller than zero (H1: µ<0). (I) Difference matrices versus the control group. The effect of delivery was tested from the difference of Z-transformed correlation coefficients between the C-section+PBS and V-delivery+PBS groups. The mean of the coefficients of this matrix was significantly smaller than zero (p<0.01, t-test). The effect size was measured using Cohen’s d (d= 0.20). The same analyses were performed on the difference between the V-delivery+IL-1β and V-delivery+PBS groups for the effect of inflammation (p<0.001, d= 0.65) and on the difference between the C-section+IL-1β and V-delivery+PBS groups for the combined effect of inflammation and delivery (p<0.001, d=0.79).

Rodents are known to communicate through ultrasonic vocalizations (USVs). Altered perinatal USV emission is a well-described phenotype associated with NDDs [47]. Pups of the C-section+PBS, V-delivery+IL-1β, and C-section+IL-1β groups emitted significantly more calls at P2 than those of the V-delivery+PBS group (p=0.046, 0.013, and 0.003, respectively; Fig. 4B). This effect was the highest in the C-section+IL-1β group.

Similarly, social impairment is often associated with NDDs [47]. We used a previously described early developmental stage social behavioral test based on nest odor affinity to assess social impairment [29, 44]. At P8, pups exposed to perinatal inflammation spent less time on the nest sawdust (p=0.014; Fig. 4C). Such disinterest at P8 was more pronounced for the C-section+IL-1β-exposed animals (p=0.0013; Fig. 4C).

In addition, we performed the three-chamber test to evaluate adolescent (6 weeks of age) social interactions (Fig. 4D) [47]. We observed that the V-delivery groups spent significantly more time in the stranger zone than in the object zone (p=0.0018 for PBS and p<0.0001 for IL-1β), indicating social interest. However, such social interest was lower in the C-section+PBS group (p=0.01) and not present in the C-section+IL-1β group (p=0.598). We obtained similar results when measuring the number of entries foreach zone: more entries into the stranger zone for the V-delivery groups, which was not observable for the C-section groups. Moreover, C-section+IL-1β animals entered the stranger zone significantly less than V-delivery+PBS animals (p=0.016, Fig.4D). The C-section+PBS and V-delivery+IL-1β groups had similar social indexes as the V-delivery+PBS group, whereas the C-section+IL-1β group had a significantly lower social index (p=0.0472, Fig.4D).

Functional ultrasound (fUS) is a recently developed imaging technique, called ultrafast Doppler, to monitor *in vivo* brain activity through fluctuations in cerebral blood volume (CBV) [48]. We averaged the relative fUS signals (which are proportional to CBV) over the pixels of each region of interest before normalization. This resulted in one signal per functional area. We then computed the pairwise correlation of those signals and arranged the correlation coefficients in a connectivity matrix. (Fig. 4E-G). This qualitative (Fig. 4F) and quantitative (Fig. 4H) analysis showed a difference in brain connectivity between PBS-exposed animals born vaginally versus those born via C-section. Specifically, the C-section+PBS group showed less brain connectivity than the V-delivery+PBS group, as indicated by lower-than-zero coefficients in the differential matrix (p<0.01; Fig. 4H-I). This suggests a global reduction in connectivity associated with C-section delivery. We made similar observations for both the V-delivery+IL-1β and C-section+IL-1β groups (p<0.001). Calculation of Cohen’s d showed the effect of C-section to be smaller (D=0.20) than that of IL-1β (D=0.65). Furthermore, fUS revealed an additive effect of C-section with IL-1β exposure, as the combined effect was more significant (D=0.79) than either condition alone.

## Discussion

The key finding of the present study is that C-section and systemic neonatal inflammation synergize to exacerbate the deleterious effects of inflammation in a mouse model of EoP. We observed such synergy for the diversity of the gut microbiota, peptidoglycan levels in plasma, microglia/macrophage reactivity, OLIG-2^+^ cell density, myelin gene expression, MBP protein expression, behavior, and, most importantly, brain connectivity.

The C-section+PBS group showed few alterations, except for diversity of the gut microbiota, the density of PDGFRα^+^ and PDGFRα^+^/O4^+^cells, the number of USV calls, and brain connectivity. One potential bias in assessing the effects of C-section+PBS was that the present study focused on parameters that we previously showed to be altered in the V-delivery+IL-1β group [16, 29–32, 44]. Other non-studied parameters could potentially have been altered in the C-section+PBS group.

Another important finding of the present study is the demonstration of increased expression of the three tested receptors for peptidoglycans on brain microglia/macrophages. This could be part of the mechanism that links the observed reduction in microbiota diversity and altered microglia/macrophage reactivity and the subsequent consequences for brain development and connectivity [38].

In conclusion, C-section and systemic inflammation are two significant risk factors for brain maldevelopment that can interact and synergize to exacerbate brain abnormalities. It the results of the present study are applicable to human preterm infants, they potentially raise the question of the risk-to-benefit balance of C-section in the context of chorioamnionitis/fusinitis/intra-uterine infection.

## Materials and Methods

### Animals and models

Experimental protocols were approved by the Ethics committee and the services of the French Ministry in Charge of Higher Education and Research (#10469). Experiments were performed on OF1 strain mice (Charles River, France). C-sections were performed on pregnant mice after cervical dislocation at E19. The surgical procedure was performed under an infrared lamp. Hysterectomy was performed after a combined transverse cutaneous incision of the abdomen and peritoneal incision. Fetuses were extracted from the uterus by manual pressure. Newly born pups were stimulated by massage and transferred to an adoptive mother at the time of the first breathing movements. Surgical procedures lasted between 3 and 5 min. V-delivery occurred spontaneously at E20. Pups from all experimental groups were immediately placed after birth with another OF1 mother that had delivered 24 h earlier.

Male pups were used for the experiments and injected intra-peritoneally (i.p.) twice a day from P1 to P5 with 10 μg/kg IL-1β (Miltenyi Biotec) diluted in 5 μL 1X PBS or with 5 μL 1X PBS alone [29, 30, 32]. Pups were monitored twice a day for any signs of distress according to a clinical score that included feeding, respiratory rate, and weight gain.

### P5 colon collection and bacterial DNA extraction and sequencing

At P5, colons were dissected (from caecum to rectum) and placed at −80°C. Bacterial DNA was extracted using a QIAamp PowerFecal DNA Kit according to the manufacturer’s protocol. The V3-V4 hyper-variable region of the 16S rRNA gene was amplified by PCR using 10 ng of fecal DNA, 0.5 μM primers (Table1), 200 µM dNTP, and 0.5 U DNA-free Taq-polymerase, MolTaq 16S DNA Polymerase (Molzym). The resulting PCR products were purified, quantified, and sequenced using Illumina MiSeq technology (Illumina, CA, USA).

### OTU table generation and statistical analysis

Sequences of the V3-V4 amplicons were processed using the FROGS pipeline to obtain abundance tables of OTUs and their taxonomic affiliation. The successive steps involved de-noising and clustering of the sequences into OTUs using SWARM, chimera removal using VSEARCH, and taxonomic affiliation for each OTU using the RDP Classifier of the SILVA SSU Pintail100 138 database. Statistical analyses were performed using “R” (v.4.0.3). α-diversity analysis was performed using the package phyloseq (v.1.34.0)[37]. For each sample, the observed species and Chao1 index were used to estimate species richness, whereas the Shannon and InvSimpson indexes were used as comprehensive indicators of species diversity and uniformity. Differentially abundant OTUs among the different groups were investigated using DESeq2 (v.1.30.0) by performing the Wald significance test with Benjamini-Hochberg false discovery rate correction. Statistical significance was set at padj < 0.05. Heatmaps were produced using Morpheus software (https://software.broadinstitute.org/morpheus), which is a hierarchical clustering method that couples means using Pearson’s correlation. The cluster annotation can be found in Table 1.

### Protein extraction and Peptidoglycan ELISA

Following decapitation at P5, blood collection was performed from the bodies and heads and then the brains were extracted. Plasma was collected after centrifugation for 10 min at 4,000 rpm. Brain proteins were extracted using RIPA Buffer (Sigma-Aldrich) containing protease inhibitors (cOmplete Tablets, Roche) in gentleMACS M tubes using a gentleMACS dissociator (Miltenyi Biotec) according to the manufacturer’s instructions. Samples were centrifuged (10,000 × g, 10 min, 4°C) and the supernatants stored for later use. ELISA using a mouse peptidoglycan (PG) ELISA Kit (MBS263268, MyBioSource) [49] was performed following the manufacturer’s instructions.

### Western Blot

Western blot analysis of MBP was performed on protein lysates from the anterior cerebrum at P22 as previously described [32] using rat anti-MBP (Millipore MAB386 1:500) and anti-β-actin (Sigma-Aldrich AC-74, 1:20,000) for staining.

### Brain dissociation and microglia/macrophage magnetic cell sorting (MACS)

P1 and P5 pups were injected with an overdose of Euthazol and perfused intracardially with 0.9% NaCl. Brains without the cerebellum and olfactory bulbs were collected and dissociated using a Neural Tissue Dissociation Kit containing papain and a gentleMACS Octo Dissociator with Heater. Magnetic beads coupled with mouse anti-CD11B antibodies (microglia/macrophage) were used for cell isolation according to the manufacturer’s protocol (Miltenyi Biotec) as previously described [29, 30, 32].

### ROS production analysis

CD11B^+^ cells (80,000) were suspended in Hank’s Balanced Salt Solution with Ca^2+^ and Mg^2+^ (HBSS^+/+^) and incubated with luminol (50 µM; Sigma) at 37°C in the dark for 10 min. Half of the cells were stimulated with phorbol 12-myristate 13-acetate (PMA, Sigma-Aldrich) to enhance basal ROS production as previously described [44, 46].

### Flow cytometry analyses (FACS)

After dissociation of the brain at P5, as already described, cells were counted using a Nucleocounter (Chemometec). The suspended cells were adjusted to a concentration of 10^7^ cells/mL by mixing with 1X Dulbeccos’s PBS containing 2 mM EDTA and 0.5% bovine serum albumin. Immunostaining was then carried out using anti-CD11b BV421, anti-CD45 BV510, and anti-CD18-APC for microglial/macrophage phenotyping and anti-O4 Vio Bright and anti-PDGFRα PE for oligodendrocyte lineage phenotyping. The samples were incubated overnight at 4°C in the dark and sample analysis performed the following day using an LSR FortessaTM X-20 device (BD Biosciences).

### RNA extraction and real-time qPCR

Brains for myelin protein gene expression were collected at P22 after cervical dislocation and placed at −80°C prior to mRNA extraction. mRNA from CD11B^+^ cells was extracted using an RNA XS Plus extraction kit and that from brains using a NucleoSpin RNA extraction kit according to the manufacturers’ protocol (Macherey-Nagel®). Reverse transcription was performed using 350 ng (CD11B^+^) and 1,000 ng (brain) mRNA with an iScript cDNA synthesis kit (BioRad). Real-time quantitative PCR was performed on triplicate samples using SYBR Green Supermix (BioRad). mRNA levels were calculated using the 2 delta Ct method after normalization against Rpl13a mRNA as the reference mRNA. Expression is expressed as that relative to vaginally born/PBS-exposed animals as previously described [32, 44, 46] (Table 2).

**Table 2.**
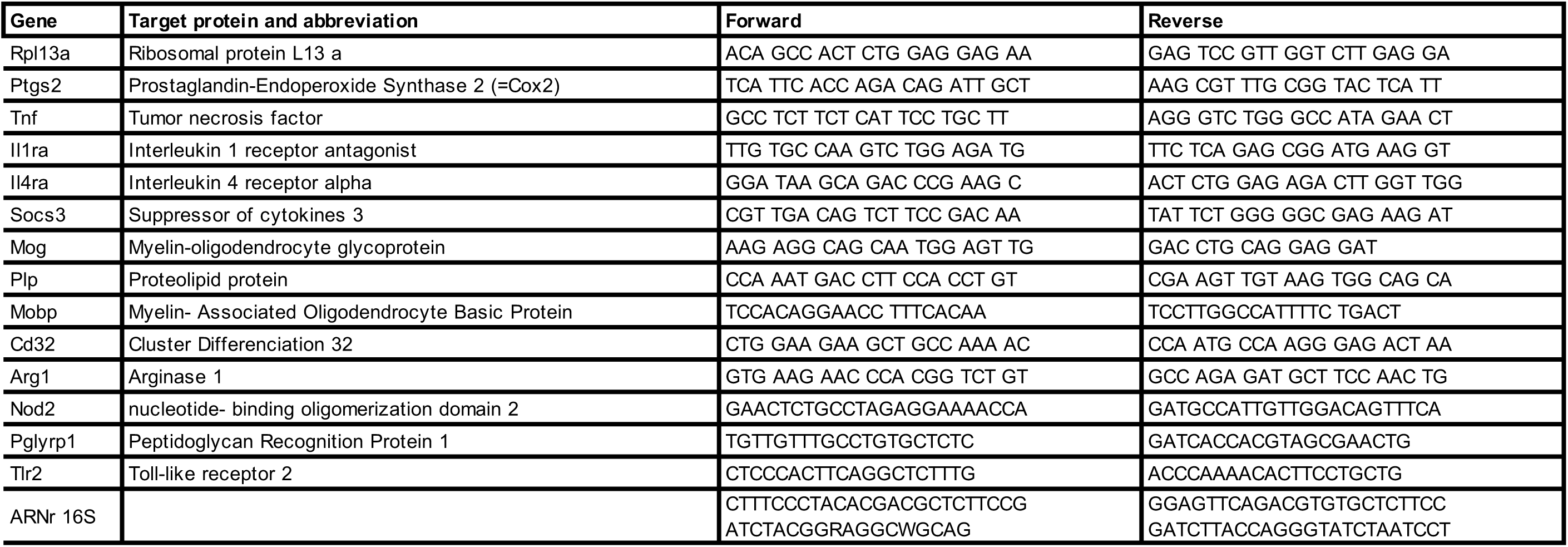

### Immunohistochemistry

At P5, brains were processed to paraffin sections by immediate immersion in 4% formaldehyde for seven days at room temperature prior to dehydration and paraffin embedding. Sections were cut using a microtome. Immunostaining was performed using a Leica Bond max robot and the BOND Polymer Refine Detection kit and mouse antibody to anti-OLIG-2 (18953, Immuno-Biological Laboratories, 1:200). Images were acquired using a Nanozoomer slide scanner (Hamamatsu) and extracted for each region of interest using QuPath software [50].

### Neonatal behavioral tests

Ultrasonic vocalizations were recorded at P2 for 3 min using an ultrasound microphone (Noldus) sensitive to frequencies from 30 to 90 kHz. The pup isolated from the litter was placed in a container (H 4.5 cm × L 10 cm × W 10 cm) inside a soundproof polystyrene box (H 23 cm × L 37 cm × W 24 cm) to avoid interference from external noise. The box temperature was 23.2 degrees. The microphone was placed 10 cm above the pup. Ultrasonic vocalizations were recorded using Ultravox XT3.1 software (Noldus) and analyzed using the https://usv.pasteur.cloud tool [51].

Social behavior at P8 was evaluated by the nest odor preference test as previously described [29]. The room temperature was 24°C. The test apparatus (L 20 cm × W 13 cm) was composed of three zones: a nest zone, with the nest, and a clean zone, with clean sawdust (L 7 cm × W 13 cm), separated by a neutral zone (L 6 cm × W 13 cm) without sawdust. The pup isolated from the litter was placed in the neutral zone at the beginning of the test and the time spent in each area measured for 1 min. The test apparatus was cleaned with 70% ethanol between pups.

### Adolescent behavioral tests

At six weeks, a three-chamber social and behavioral test was conducted on the mice. The experiment involved placing the mouse in a three-chamber box with openings between the chambers. The test was performed in two sessions. During the first session, the mouse was allowed to freely explore the empty apparatus for 10 min. The following day, the mouse was placed in the middle chamber while a stranger mouse was placed in one of the chambers (either left or right) and an object placed in the other chamber. The mouse being tested was allowed to freely move around the entire apparatus for 10 min. The experiments were recorded and analyzed using Any-Maze software. The analysis was based on the time spent in each chamber and the number of entries made. The social Index was calculated as follows:

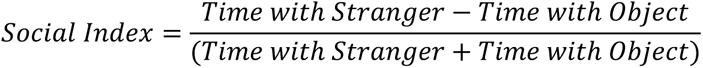

### Ultrafast Doppler acquisition

At six weeks, animals (N=38) were anesthetized with a blend of ketamine (75 mg/kg) and xylazine (15 mg/kg) per animal. A craniotomy was performed after fixing their head in a stereotaxic frame.

Functional ultrasound images were acquired at a frame rate of 2.5 Hz using an Iconeus One ultrasound system (Iconeus, Paris, France) with an IcoPrime linear ultrasound probe (15 MHz, 128 elements, pitch 0.11mm). The probe was placed at the coronal plane showing the somatosensory cortex, thalamus, hypothalamus, and hippocampus. Detailed explanations for atlas registration and brain positioning can be found in [52, 53]. fUS images were recorded for 15 min with a field-of-view that was 14 mm wide and 11 mm deep. Recordings from animals presenting with a hemorrhage, stroke, or spreading depression were excluded from the analysis. Epochs with motion or other artifacts were also removed for each animal during data processing. Finally, the fUS signal for each pixel (proportional to CBV) was filtered using a bandpass filter (4^th^ order Butterworth) with frequency cut-offs of 0.01 and 0.1 Hz and transformed as a relative value (%) to the average.

We calculated the mean matrices for each animal to compare connectivity patterns between conditions. To ensure that the values were normally distributed after transformation, we performed a Z-Fisher transformation of the correlation coefficient. Then, we subtracted the mean matrix of the control group (vaginally born/PBS-exposed animals) from the mean matrix of each other condition. We hypothesized that the “difference coefficients” would be, on average, negative, which we tested using a one-tail t-test on the coefficient of the matrices of differences (H1 hypothesis: difference coefficient < 0). For a more quantitative understanding of the modification of connectivity of each condition, we computed Cohen’s d, which measures the effect size (typical values: d = 0.2 means small effect, d = 0.5 medium effect, d = 0.8 large effect).

### Statistical analysis

Data are expressed as mean values with the standard error of the mean (SEM). GraphPad Prism Software was used for all statistical analysis. Multiple comparisons for the same data set were performed by one-way analysis of variance (ANOVA) using the Kruskal-Wallis test followed by an uncorrected Dunn’s test or two-way ANOVA followed by an uncorrected Fisher’s LSD test in comparison to the V-delivery+PBS group.

## Acknowledgments

Schematic representations were created using BioRender.com. PG acknowledges support from Inserm, the Université Paris Cité, Fondation Grace de Monaco, Inserm International Research Project “IntegrA”, and French “Investissement d’Avenir-ANR-11-INBS-0011 NeurATRIS”. JVS acknowledges support from the Agence Nationale de la Recherche ANR-22-CE14-0051-02 and the Fondation de France. CB acknowledges support from “Investissement d’Avenir-ANR-17-EURE-001-EUR G.E.N.E.’’ and the ANR (contract ANR-22-CE37-0019). BM acknowledges support from Inserm, the Université Paris Cité, Fondation Air Liquide, Fédération pour la Recherche sur le Cerveau, and the Région Ile-de-France. RDH acknowledges support from the Swedish Research Council (2018-06232). BF acknowledges support from the Cerebral Palsy Alliance, Australia. SA acknowledges support from INRAE. ADD acknowledges support from the Université Sorbonne Paris Nord and the French National Research Agency (#ANR-18-CE17-0009-01, #ANR-18-CE37-0002-03, #ANR-21-RHUS-009). MT acknowledges support from the Inserm Accelerator of Technological Research on Biomedical Ultrasound. PG, FF, BF, MT, and CB acknowledge support from the European Union’s Horizon 2020 Research and Innovation program under Grant Agreement No 874721 PREMSTEM. The supporting bodies played no role in any aspect of study design, analysis, interpretation, or the decision to publish this data.

---

**Supplementary Figure 1.**
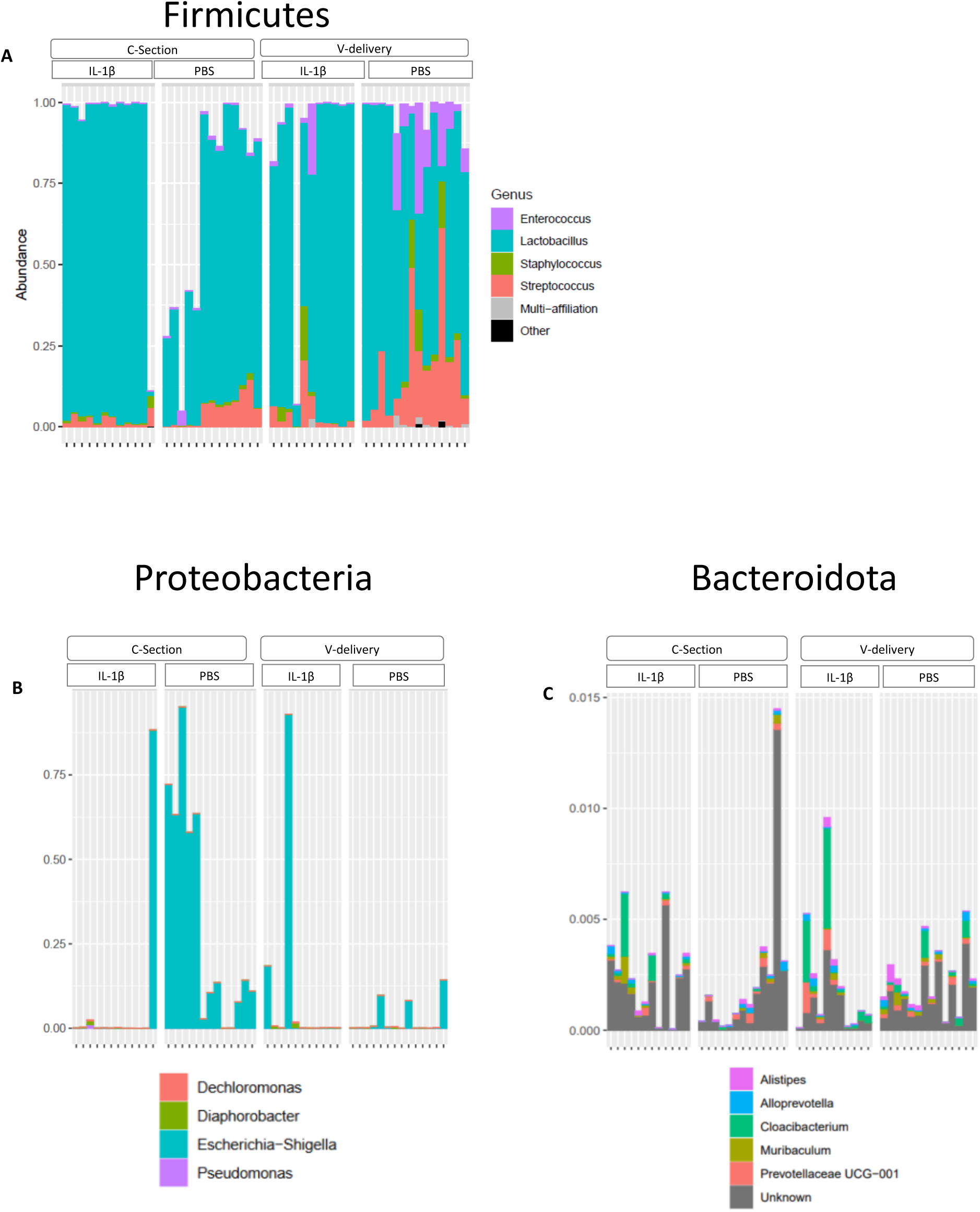
Impact of IL-1β and C-section on microbiota composition per phylum. Table of the abundance of operational taxonomic units (OTUs). Analysis of the individual distribution of the Firmicute (A), Proteobacteria (B), and (Bacteroidota) composing the intestinal microbiota at P5 in V-delivery ± IL-1β and C-section ± IL-1β mice.

**Supplementary Figure 2.**
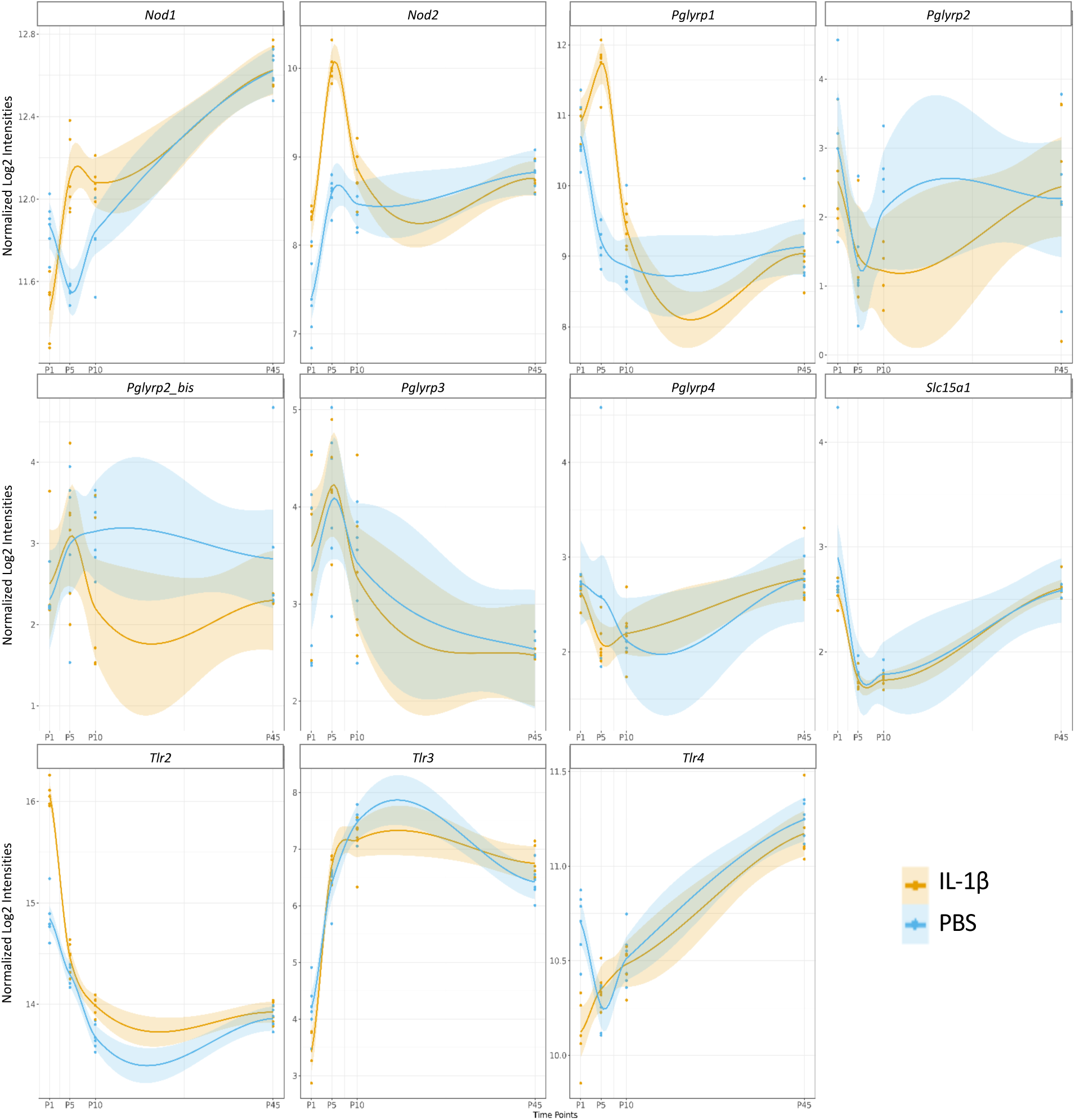
Impact of IL-1β on microglial expression of peptidoglycan receptors. Relative expression of microglial peptidoglycan receptor mRNA over time in CD11B^+^ cells of V-delivery ± IL-1β mice. The datasets came from Krishnan et al. [31] and Van Steenwinckel et al. [32].

